# A globally synthesised and flagged bee occurrence dataset and cleaning workflow

**DOI:** 10.1101/2023.06.30.547152

**Authors:** James B. Dorey, Erica E. Fischer, Paige R. Chesshire, Angela Nava-Bolaños, Robert L. O’Reilly, Silas Bossert, Shannon M. Collins, Elinor M. Lichtenberg, Erika M. Tucker, Allan Smith-Pardo, Armando Falcon-Brindis, Diego A. Guevara, Bruno Ribeiro, Diego de Pedro, John Pickering, Keng-Lou James Hung, Katherine A. Parys, Lindsie M. McCabe, Matthew S. Rogan, Robert L. Minckley, Santiago J.E. Velazco, Terry Griswold, Tracy A. Zarrillo, Walter Jetz, Yanina V. Sica, Michael C. Orr, Laura Melissa Guzman, John S. Ascher, Alice C. Hughes, Neil S. Cobb

## Abstract

Species occurrence data are foundational for research, conservation, and science communication, but the limited availability and accessibility of reliable data represents a major obstacle, particularly for insects, which face mounting pressures. We present *BeeBDC*, a new *R* package, and a global bee occurrence dataset to address this issue. We combined >18.3 million bee occurrence records from multiple public repositories (GBIF, SCAN, iDigBio, USGS, ALA) and smaller datasets, then standardised, flagged, deduplicated, and cleaned the data using the reproducible *BeeBDC R*-workflow. Specifically, we harmonised species names (following established global taxonomy), country names, and collection dates and we added record-level flags for a series of potential quality issues. These data are provided in two formats, “cleaned” and “flagged-but-uncleaned”. The *BeeBDC* package with online documentation provides end users the ability to modify filtering parameters to address their research questions. By publishing reproducible *R* workflows and globally cleaned datasets, we can increase the accessibility and reliability of downstream analyses. This workflow can be implemented for other taxa to support research and conservation.

## Background & Summary

Occurrence datasets are increasingly critical for scientific research, conservation, and communication worldwide. From foundational systematic^1,2^, life-history^3^, and conservation studies^4,5^, to continental or global macroecological^6,7^ or macroevolutionary analyses^8^, occurrence data are used to understand the natural world and form the basis of research, policy, and management. Spatiotemporal occurrence records (e.g., from specimens, images, or observations) are being delivered to, and produced by, community scientists in quantities that were unthinkable just a decade ago, with ever-increasing identification support by professional experts^9^. However, community-generated records can be biased towards larger and more charismatic taxa^10^. Online repositories, such as the Global Biodiversity Information Facility (GBIF) and Symbiota Collection of Arthropod Network (SCAN), are invaluable data aggregators and tools in supporting the mobilisation of arthropod occurrence data to help both researchers and general audiences understand biodiversity. Concurrently and complementarily, initiatives like iNaturalist and QuestaGame (Australia) enable community and professional scientists to generate occurrence data for research, science communication, and more^9^. Occurrence data are being generated daily, making new analyses possible for previously data-poor taxa. These data are key to developing National Strategic Biodiversity and Action plans, upon which the monitoring framework of the Post-2020 Global Biodiversity Framework will depend^11^. Therefore, developing methods to aggregate and standardise such data is essential.

Despite the great utility of occurrence data, many issues can inhibit their successful use. Firstly, whilst “open data” is now mandated in many journals, data standards are inconsistently implemented. This means that these data are increasingly fragmented and thus still inaccessible. Occurrence records can be rendered partially or critically incomplete at many different phases between the initial point of data collection and point of data publication and sharing^12^. For example, poor data collection practices at the point of production (in the field) can make downstream recovery efforts impossible. Data can often be omitted or altered during transcription, digitisation, or geo-referencing, making occurrences unusable or misleading. For example, only 2.7 out of 31 million (9%) Brazilian plant occurrences were considered high-quality following data cleaning and validation by Ribeiro, et al. ^13^. Misidentification can also be a major issue; for example, 58% of African gingers from 40 herbaria across 21 countries bear an incorrect name^14^, highlighting the importance of accessible museum and herbarium specimens^15^. Secondly, data can be of excellent quality, digitised, and used in publications, but they may not be made available in accessible repositories. Whatever the reason, such data are effectively unavailable for — or, in the case of misleading data, detrimental to — further research, conservation, and management^16^. Funding institutions to curate and digitise collections might fill in some of the major data gaps revealed in recent global analyses of public data^6^.

Mobilising massive datasets with numerous potential issues represents a major roadblock for researchers and other users. A careful balance between discarding too much data and accepting too many problematic records must be maintained. Ideally, problematic records can instead be flagged (allowing users the choice of treatment) or fixed using a combination of automated pipelines and expert validation^17^. The barriers are such that the data cleaning process is often built from scratch or nearly so for each new project (re-inventing the wheel)^6,7^. Constantly rebuilding the data cleaning process can result in haphazard application of cleaning and data quality and is a major hurdle for many researchers and conservation practitioners that might preclude them from undertaking robust and reproducible analyses. This represents a major accessibility problem that stalls research, management, and conservation — particularly for research groups that lack sufficient support, expertise, facilities, or time to develop or implement cleaning workflows — that could otherwise produce excellent research. This results in major knowledge gaps of species distributions and ecological niches, especially in developing economies and over extended timescales^18^. Until now, very few major efforts have been made to combine, clean/flag, and make accessible the world’s occurrence datasets; this is particularly true for diverse and ecologically important arthropods^7,19,20^. We tackle this issue of data cleaning and reproducibility using a major group that is taxonomically extensive (>20,000 valid species), ecologically and economically important, and for which research is booming globally: bees.

Bees (Hymenoptera: Anthophila) are a keystone taxon of terrestrial ecosystems across the globe and the most significant group of pollinating animals in both agricultural and natural settings^21,22^. As such, great efforts are made to understand their spatial distributions^6^ and it is critical for research on this flagship invertebrate taxon to be robust and correctly analysed. Despite the urgent need for quality occurrence data, there are many barriers to their use, particularly accessibility and reliability. Removing these hurdles will encourage new research and enable best practices at greatly reduced effort and financial cost. At the time of download and to our knowledge, there are ~18.1 million bee occurrence records that are publicly available in major public repositories and more that are private or otherwise inaccessible. Here we collate and process several bee datasets for open-use by all interested parties. We also produce a machine-readable version of the global bee taxonomy and country-level checklist available on the Discover Life website^23^. Specifically, we curated multiple raw datasets, combined curated records, taxonomically harmonized species names, and annotated potentially problematic records. Herein, we combined public and private datasets to provide (i) a cleaned and standardized dataset (6.9 million occurrences), (ii) a flagged-but-uncleaned dataset (for users to filter based on our flagging columns; 18.3 million occurrences), (iii) all R-scripts and input files needed to produce both datasets (Fig. 1), and (iv) summary figures and tables highlighting the state of global bee data. All data and scripts are accessible, documented, and open access. Importantly, we plan to periodically update and publish new versions of the *BeeBDC* occurrence data cleaning package^24^ and bee data. This will leave room for improved versions and functionality, especially because new occurrence data are consistently being added to online databases. Despite being developed specifically for bee datasets, many of our functions are generic and can be applied to various taxa for which similar issues exist.

**Figure 1.**
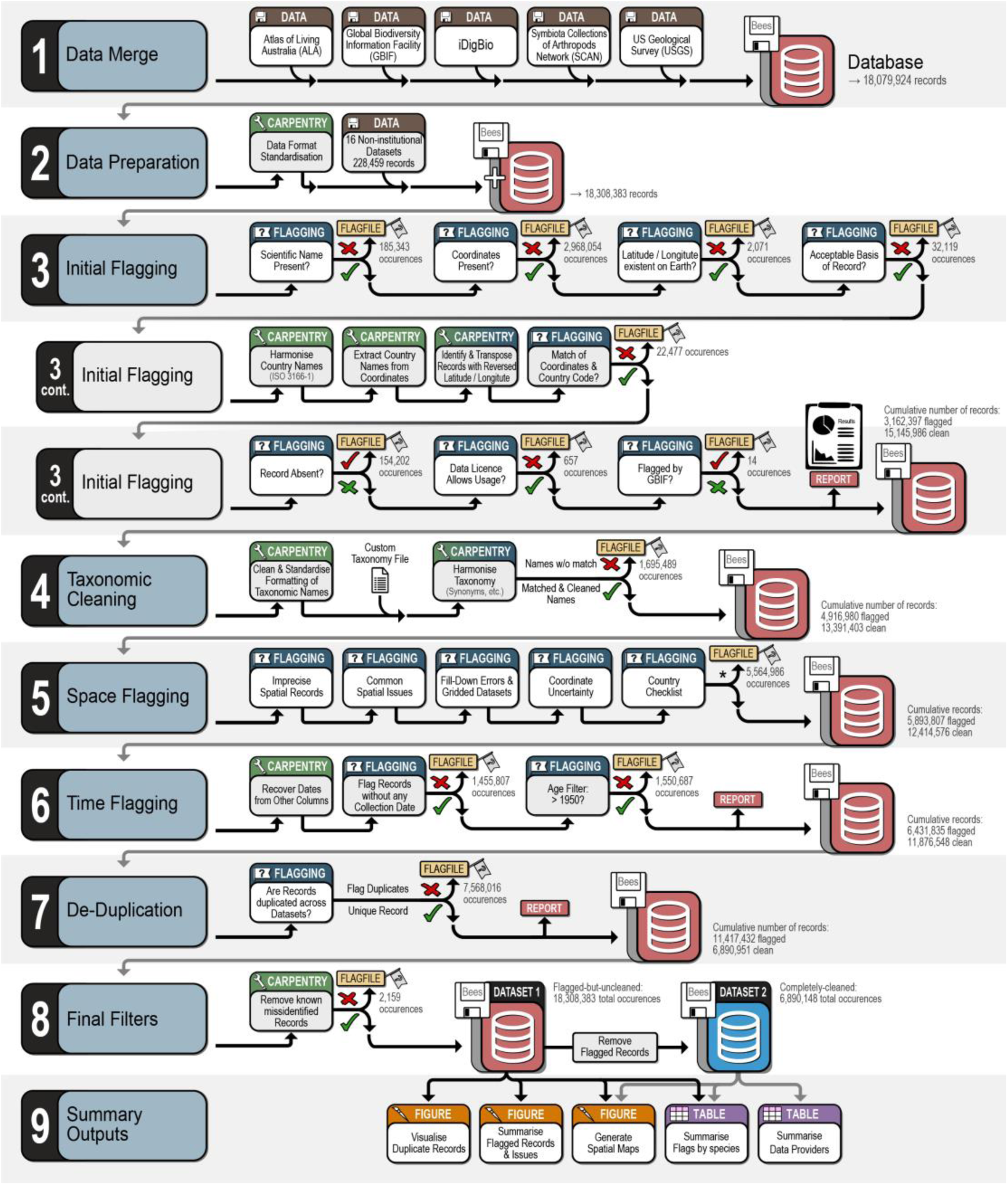
Visual summary of our BeeBDC^24^ workflow for compiling, cleaning, flagging, and summarising the most extensive, publicly available data set of native bee occurrences. Asterisk indicates that the number of records flagged in (5) Space Flagging includes all steps.

## Methods

### Data sources

We sourced data from publicly available online data repositories (https://doi.org/10.25451/flinders.21709757; ExtraTables/MajorRepoAttributes_2023-09-01.xlsx)^25^ as well as select non-public, private, or otherwise publicly inaccessible sources that were willing to share their data for this study. We use the term “data source” in the manuscript, but recognise that these sources can be very distinct in function. For example, aggregators serve data that is created and curated by multiple distinct data providers. The primary repositories that we sourced data from were (i) GBIF, (ii) SCAN, (iii) Integrated Digitized Biocollections (iDigBio), (iv) the United States Geological Survey (USGS), and (v) the Atlas of Living Australia (ALA). Additionally, we have sought out and incorporated smaller public or private data sources from (i) Allan Smith-Pardo (ASP), (ii) Robert Minckley (BMin), (iii) Elle Pollination Ecology Lab (EPEL^26^), (iv) *Bombus* Montana (Bmont; Casey Delphia^27^), (v) Ecdysis (Ecd^28^), (vi) Gaiarsa, et al. ^29^ (Gai), (vii) the Connecticut Agricultural Experiment Station (CAES^30,31^), (viii), USDA ARS South-eastern USA (Parys), (ix) Eastern Colorado (Arathi Seshadri), (xi) Florida State Collection of Arthropods (FSCA), (xi) Armando Falcon-Brindis (Arm), and (xii) four more publicly available bee datasets from the literature (SMC, Bal, Lic, Dor, VicWam)^3–5,32–39^. These datasets were collated by directly engaging and collaborating with their owners, particularly where data gaps were perceived, but with a particular emphasis on the Americas.

Global Biodiversity Information Facility data were downloaded via the online portal on the 14^th^ of August 2023^40–46^ on a per-family basis. SCAN data were downloaded between the 14^th^ and 20^th^ of August 2023^47–53^ on a per-family basis and by subsets of collections if the one million record limit was surpassed (the maximum capacity for a single download). Integrated Digitized Biocollections data were downloaded on the 1^st^ of September 2023^54–60^ by selecting all Apoidea records and filtering to the seven bee families (Andrenidae, Apidae, Colletidae, Halictidae, Megachilidae, Melittidae, and Stenotritidae). United States Geological Survey data were provided directly by Sam Droege on the 19^th^ of November 2022^61^. Finally, ALA data were downloaded on the 1^st^ of September 2023^62^ using a *BeeBDC* wrapper function, *atlasDownloader*, which uses their *R*-package, *galah*^63^.

We sourced taxonomic name and checklist data from information hosted on the Discover Life website^23^. Our bee taxonomy is current as of the 20^th^ of August 2023 and the bee country checklist as of the 21^st^ of August 2023. Both of these datasets are maintained by Ascher and Pickering ^23^ and are critical sources to ensure occurrence quality. We also added small updates to both datasets based on those published in Orr, et al. ^6^ and to reflect orthographic variants that were identified during manual data and error checks. We undertook manual checks of over 4,500 flagged interactive bee species maps (mostly in the Americas) to highlight and correct potential errors in these datasets and our functions.

### Data-cleaning workflow

Analyses were undertaken in *R* version 4.3.1^64^ and *R-Studio* “Beagle Scouts” Release. The *BeeBDC* package and script relied heavily on several *R-*packages. Much of the workflow script used and built upon the *bdc* (biodiversity data cleaner) package^13^, tidyverse packages — particularly *dplyr*^65^, *magrittr*^66^, *tibble*^67^, *stringr*^68^, *tidyselect*^69^, *ggplot2*^70^, *tidyr*^71^, *rlang*^72^, *xml2*^73^, *readr*^74^, and *lubridate*^75^, — and *CoordinateCleaner*^76^. Additionally, we used *R.utils*^77^, *galah*^63^, *emld*^78^, openxlsx^79^, *rnaturalearth*^80^, *rnaturalearthdata*^81^, *countrycode*^82^, *hexbin*^83^, *cowplot*^84^, *ggspatial*^85^, *renv*^86^, *chorddiag*^87^, *igraph*^88^, *sf*^89^, and *terra*^90^. In some cases we have modified or rebuilt *bdc* functions into *BeeBDC* to better work with other parts of our package or to produce different results. Functions that relate strongly to *bdc* are prefixed with “jbd_” to indicate to users that an alternate function exists and to compare package documentation (e.g., by running “?*jbd_coordinates_precision*” in *R*) for each. Our script was both built and run on a 2019 MacBook Pro with a 2.4 GHz 8-core Intel i9 with 64GB 2667 MHz DDR4 RAM. Hence, while it can be run on personal computers — and this accessibility is intentional — we are aware that memory (RAM) and physical storage could inhibit some devices from processing the full dataset (i.e., 18.3 million occurrences and ~132 columns).

The core workflow, *R* vignette, and functions are freely available on GitHub (https://jbdorey.github.io/BeeBDC/index.html). The workflow is clearly numbered and labelled using the *R-Studio* document outline to allow quick navigation of the script. For clarity and continuity, we list the sections below according to our script, including (0.x) script preparation, (1.x) data merge, (2.x) data preparation, (3.x) initial flags, (4.x) taxonomy, (5.x) space, (6.x) time, (7.x) de-duplication, (8.x) data filtering, (9.x) summary figures and tables, and (10.x) package data. Additionally, most of our functions attempt to provide extensive and informative user-outputs for quality assurance. We provide a reference table of the occurrence-cleaning functions that are available between *BeeBDC*, *bdc*, and *CoordinateCleaner* (Table S1). Our script focuses on the specifics of bee data; however, it provides the templates needed to integrate the specifics of other taxon groups and input data.

Most functions in our script aim to identify and flag potentially problematic occurrence records based on specific tests and user-provided thresholds (flagging functions). However, several functions modify the occurrence records themselves when errors are identified (carpentry functions). We also provide summary and filtering functions that allow users to explore data issues and export useable datasets. While most functions can be implemented anywhere in the workflow, a subset relies on columns produced from earlier functions and these are highlighted in the package documentation and website. Below, we explain the sections and functions found in our script and the logic behind their implementation. We do not explain the optional steps documented in the script as they were not implemented in our dataset, but see the *BeeBDC* documentation for further clarification.

### 0. Script preparation

To ensure accessibility and ease of use, we ask users to set the root file path in which the script should look for or create certain datasets or functions. Our *dirMaker* function should locate and, if missing, create the file structure sought by the rest of the script and the *bdc* package. Packages are then installed and loaded. Here the script initialises and stores the *renv* files. The *renv* package files keep information about package versions to encourage further reproducibility of scripts and ease of publishing for users.

### 1. Data merge

The data merge section of the script reads very large datasets from the major data repositories — GBIF, iDigBio, USGS, SCAN, and ALA. It then (i) unifies the data (selecting certain columns set by the *ColTypeR* function) to Darwin Core format^91^, (ii) merges them into a single dataset, and (iii) creates a metadata file that both accompanies the *R*-object and is saved locally (ExtraTables/MajorRepoAttributes_2023-09-01.xlsx)^25^. These functions were built to accommodate the entirety of the large repositories above; however, they can be slow on computers with limited RAM.

### 2. Data preparation

Here, we provide users with two options to re-import the data saved during *1. Data merge*. For general applications, the *bdc* package provides excellent import functionality but would require manual work initially (2.1a in script). For users who want to run data from the above major repositories, we provide seamless functionality to read those data (2.1b in script). For users aiming to update the bee datasets from online repositories, they may incorporate the documented manual data edits (2.2 in script) and re-incorporate several privately curated datasets (2.2–2.5 in script). Once a dataset is standardised to Darwin Core format and contains the necessary columns, users can begin flagging or cleaning each occurrence record for data integrity. Our total dataset comprised 18,308,383 uncleaned bee occurrence records (Fig. 1).

### 3. Initial flags

To perform initial data-quality flags and checks, we used a combination of *BeeBDC* and *bdc* functions, which we will list below. These include data flagging and carpentry functions, with the latter supporting the former.

#### 3.1 Scientific name flags

We used the *bdc* function, *bdc_scientificName_empty,* to flag records with no scientific name provided (i.e., an empty *scientificName* cell). This step flagged 185,343 (1%) occurrences.

#### 3.2 Missing Coordinates

We used the *bdc* function, *bdc_coordinates_empty,* to flag records with no coordinates provided (empty *decimalLatitude* or *decimalLongitude* columns). This step flagged 2,968,054 (16%) occurrences.

#### 3.3 Coordinates out of range

We used the *bdc* function, *bdc_coordinates_outOfRange,* to flag records that were not on the map (i.e., not between −90 and 90 for latitude or not between −180 and 180 for longitude). This step flagged 2,071 (<1%) occurrences.

#### 3.4 Poor record source

We used the *bdc* function, *bdc_basisOfRecords_notStandard,* to flag occurrences that did not meet our criteria regarding their basis of records. Broadly, we kept all event, human observation, living specimen, material/preserved specimen, occurrence, and literature data. For this step, we were relatively liberal in what we allowed to remain in the dataset and only excluded fossil specimens; however, some users might prefer to also remove human observations. This step flagged 32,119 (<1%) occurrences.

#### 3.5 Country name

We used our function, *countryNameCleanR*, to match GBIF ISO2 country codes to country names from a static Wikipedia ISO2 to country name table within the function. Users can also input a data frame of problem names and fixed names to replace them in the dataset. We then created a modified version of the *bdc* function *bdc_country_from_coordinates* into a chunking function, *jbd_CfC_chunker* (to work on smaller portions of the dataset at a time), where the user can specify chunk-sizes to best manage this RAM-intensive function. This function also can be run in parallel (multiple threads). However, for all parallel-ready functions users must be aware of their available RAM; ten cores seemed reasonable on a 64 GB machine (mc.cores = 10) for this function, but for the remaining functions we used between two and six. The *jbd_CfC_chunker* function assigns country names to those occurrences that were missing them but that have valid coordinates that correspond to a country. This step assigned country names to 323,858 (2%) occurrences.

#### 3.6 Standardise country names

We used the *bdc* function, *bdc_country_standardized,* to attempt to further standardise country names for consistency. This step standardised country names for 4,509,749 (25%) occurrences.

#### 3.7 Transposed coordinates

We used a parallel-ready (mc.cores = 4) chunking function, *jbd_Ctrans_chunker*, (as *jbd_CfC_chunker* above) that wraps a custom version of the *bdc* function, *bdc_coordinates_transposed* (*jbd_coordinates_transposed*), which flags and corrects coordinates for which the latitude and longitude are transposed. This step corrected transposed coordinates for 2,267 (<1%) occurrences.

#### 3.8 Coordinates and country inconsistent

Here, we created a similar, but parallel-ready (mc.cores = 4), function to *bdc*’s called *jbd_coordCountryInconsistent*. We re-built the *bdc* version for memory efficiency and to accommodate the processing of our dataset, which is much larger than the *bdc* test dataset. Our function flags occurrences for which the coordinates and country do not match while allowing for a user input map buffer (in decimal degrees). This step flagged 22,477 (<1%) occurrences.

#### 3.9 Georeference issue

We ran the *bdc* function, *bdc_coordinates_from_locality,* which highlights occurrences for which there is no latitude or longitude directly associated with the data, but for which sufficient locality data may be used to extrapolate relatively accurate geological coordinates. This step flagged 2,388,168 (13%) occurrences.

#### 3.10 Absent records

We provide a function, *flagAbsent*, to flag occurrence records that are marked as “ABSENT”. Users may not be aware that many thousands of records are often provided as absent. These records might occur, for example, due to repeated sampling programmes not finding a certain species on a particular date or at a particular site. This step flagged 154,202 (1%) occurrences.

#### 3.11 Restricted licenses

We provide a function, *flagLicense*, that aims to flag records that are not licensed for use. While such records should not be on public data repositories, some few are. Therefore, users should be aware not to use these protected data points. This step flagged 657 (<1%) occurrences.

#### 3.12 GBIF issues

We provide a function, *GBIFissues*, to flag occurrences already flagged for user-specified GBIF issues. For example, we flagged GBIF occurrences marked as “COORDINATE_INVALID” and “ZERO_COORDINATE”. This step flagged 14 (<1%) occurrences.

At the end of the initial flags section, we provide the *flagRecorder* function to save the flags for all occurrences as a separate file. Users may then use *bdc* functions to create summaries, reports, and some figures.

### 4. Taxonomic cleaning

While *bdc* provides great functionality for taxonomic cleaning of some groups, it does not provide access to the most-current bee taxonomy. Hence, we used the online global bee taxonomy available through Discover Life^23^, which lists approximately 21,000 valid names and 31,000 synonyms or orthographic variants. Discover Life is the leading source of bee names and is peer-reviewed by global experts on an ongoing basis. We flagged this list for ambiguous names (homonyms) that, to varying degrees, could not be matched due to their complicated taxonomic histories. Records with ambiguous names were either processed differently or excluded from the below function and hence the final occurrence dataset. We also made some manual corrections as taxonomic issues (mostly orthographic variants) were identified (ExtraTables/AddedTaxonomyVariants.xlsx)^25^. The final taxonomy is available in the *BeeBDC* package as *beesTaxonomy*.

We first cleaned our species names using the *bdc* function *bdc_clean_names,* which attempts to clean names and unify writing-styles (e.g., capitalization and punctuation). It also removes family names, qualifiers (this is flagged), infraspecific terms, separates authors, dates, and annotations. This function also adds a taxonomic uncertainty flag, .*uncer_terms*, which flagged 97,231 (<1%) records. We then used our parallel-ready (mc.cores = 4) *harmoniseR* function to harmonise occurrence names with our combined Discover Life taxonomy. This function takes steps to (i) compare occurrence names directly with valid names (genus species authority), (ii) combine the *bdc*-cleaned name and occurrence authority to match against the valid name, (iii) match species names with canonical flagged names (i.e., scientific names with flags from Discover Life), (iv) directly compare scientific names, (v) combine the occurrence scientific name and occurrence authority to match against the valid name, (vi) match occurrence’s scientific name with the taxonomy’s valid name with subgenus removed from both, and (vii) as above but using canonical names. Between each step, the function filters out already-matched names. These steps are first taken for the non-ambiguous names (according to our taxonomy) and then applied again for the ambiguous names but also matching the authority (“author year”, all lowercase and with no punctuation). If a name cannot be unambiguously matched at any level (homonyms), then they will fail this function and an accepted name will not be applied. The function then moves the provided name to *the verbatimScientificName column and updates the scientificName, species, family, subfamily, genus, subgenus, specificEpithet, infraspecificEpithet, and scientificNameAuthorship* columns where better matches are found. This is only done for occurrences with a successful match; unmatched occurrences are not altered. Occurrence records that do not match the taxonomies are flagged accordingly with the *.invalidName* column. The data produced from each step are merged and an updated object is output. The minimum requirement for this function is a data frame with a column containing species names — this is intended to allow quick checking and updating of simple species lists. This step matched valid names to 15,799,107 (86%) and flagged 2,509,276 (13%) occurrences, respectively.

### 5. Space flagging

We further flagged our data for spatial issues using a combination of functions from *BeeBDC, bdc*, and *CoordinateCleaner*. We outline these steps below.

#### 5.1 Coordinate precision

We used our function, *jbd_coordinates_precision*, to flag occurrence records with latitude and longitudes below a threshold of two decimal places (~1.1 km at the equator). This step differs from the *bdc* function *bdc_coordinates_precision* by only flagging occurrences where both latitude *and* longitude were rounded. This step flagged 3,649,158 (20%) occurrences.

#### 5.2 Common spatial issues

We then used the *CoordinateCleaner* function, *clean*_*coordinates,* which runs several tests to flag potentially erroneous data. We flagged records that were within (i) 1 km of capital cities, (ii) 500 m of province or country centroids, (iii) 1 km of GBIF headquarters in Copenhagen, Denmark, or (iv) 100 m of biodiversity institutions; or that have (v) equal latitude and longitude coordinates, or (vi) zero as latitude and longitude. For example, these issues can arise when occurrences were labelled only with the city, province, country, institution, or repository are georeferenced to those exact locations. The latter two could arise from copy errors or incomplete data. We did not flag points in the ocean as they are flagged with a small buffer by *5.5 Country Checklist* (below). This step flagged (i) 15,653; (ii) 17,494; (iii) 11; (iv) 80,558; (v) 11,083; and (vi) 10,932 (each <1%) occurrences, respective to the above criteria.

#### 5.3 Fill-down errors and gridded datasets

We provide the parallel-ready (mc.cores = 4) *diagonAlley* function that uses a sliding window to flag potential fill-down errors in the latitude and longitude columns. The function removes any empty values and then groups data by event date and collector. It then arranges latitude and longitude, removes identical values, and flags sequences of latitudes or longitudes where the differences between records are exactly equal. The user defines a minimum number of repeats required for a flag (minRepeats = 6) and a minimum number of decimal places required to consider an occurrence (using 5.1 *jbd_coordinates_precision*; ndec = 3). Secondly, we then implemented the *cd_round* function from *CoordinateCleaner* to identify datasets (using the *datasetName* column) that have their latitudes or longitudes potentially gridded, using the default values. This step flagged 390,747 (2%) and 113,617 (<1%) occurrences for fill-down and gridded datasets, respectively.

#### 5.4 Coordinate uncertainty

We provide the *coordUncerFlagR* function that flags records by a user-defined threshold for coordinate uncertainty — usually provided in the Darwin Core *coordinateUncertaintyInMeters* column. We used a threshold of 1 km. This step flagged 2,831,456 (15%) occurrences.

#### 5.5 Country Checklist

We provide the parallel-ready (mc.cores = 4) *countryOutlieRs* function that uses the country-level checklist available on the Discover Life website (beesTaxonomy). While vagrants are only a minor issue, Discover Life excludes port vagrants and clear misidentifications; however, “natural” and established vagrants are generally included as valid. For example, the Fiji checklist includes eight relatively recent, but established, introductions^23,92^. The function checks all of the harmonised species names against the checklist and the country in which the occurrence falls (using overlap with *rnaturalearth*). Points that don’t align with *rnaturalearth* (e.g., they are on the coast) can be buffered by a user-specified amount, in degrees, to attempt a match. In our case, we used 0.05 degrees (~5.6 km). It is worth noting that changing the *rnaturalearth* resolution could lead to slight variations in results and that the higher resolutions (scale = 50 or “large”) might be optimal for most regions; especially for discontinuous island groups. The function produces three columns. The first column, *countryMatch*, summarises the occurrence-level result: where the species is not known to occur in that country (noMatch), it is known from a bordering country (neighbour), or it is known to occur in that country (exact). The second column’s output depends on user input. If the user wants to keep occurrences that are either exact matches or in adjacent (bordering) countries to those in the checklist (keepAdjacentCountry = TRUE) then the filtering column .*countryOutlier* will be TRUE for these cases and FALSE only for those with noMatch. A keepAdjacentCountry = FALSE argument will only flag exact country checklist matches as TRUE. For our dataset, we chose the former. The third column, .*sea*, flags the points which don’t align with *rnaturalearth* or its buffer — they are identified as being in the ocean. The *countryOutlier* column flagged 1,679,298 (9%) occurrences and the .*sea* column flagged 204,030 (1%).

Users may then produce maps, reports, and figures using the *bdc* package. They can also append the saved flag file using the *flagRecorder* function.

### 6. Time flagging

The next major sequence of functions aims to recover or filter occurrence records with date-related issues. We outline the steps taken below.

#### 6.1 Recover missing event date

We provide the *dateFindR* function that seeks to recover occurrence records that would otherwise be removed due to missing event date. The function first removes unreasonable dates based on a user-defined year-range (we removed dates prior to 1700 and after the present year). The function then seeks to harmonize date formats in a step-wise manner by first (i) combining dates from the year, month, and day columns into the *eventDate* column, then (ii) taking just the year column where it is provided without event date. Subsequently, the function looks for dates in the *verbatimEventDate*, *fieldNotes*, and *locationRemarks* columns by looking in sequence for unambiguous date strings in the following formats: (iii) year-month-day, (iv) day-month-year (where day is identifiable — i.e., days >12; or months are identifiable — in written or Roman-numeral formats), (v) month-day-year (where days or months are identifiable as above), and (vi) month-year (where month is identifiable as written or Roman-numeral formats). The function then finds ambiguous date strings in the *verbatimEventDate*, *fieldNotes*, and *locationRemarks* columns and keeps only the year value. All date values are then returned in the standardised year-month-day-hours-minutes-seconds format. Users should consider this threshold critically in relation to their own hypotheses and potentially run *dateFindR* over the Flagged-but-uncleaned dataset with their own threshold if required. This step rescued dates for 1,333,087 (7%) occurrences.

#### 6.2 Missing event date

We then used the *bdc* function, *bdc_eventDate_empty*, to flag records that do not have date in the *eventDate* column. This step flagged 1,455,807 (8%) occurrences.

#### 6.3 Old records

We used the *bdc* function, *bdc_year_outOfRange*, to flag occurrence records that are likely too old for use in species distribution modelling. Here, we flagged occurrences from before 1950. Users may want to consider changing this value or disregarding this flag as it applies, or does not, to the question(s) that they want to ask. This step flagged 1,550,687 (8%) occurrences.

Users may then create reports and figures on the time filters using the *bdc* package. They can also append the saved flag file using the *flagRecorder* function.

### 7. Duplicate records

Duplicate records frequently arise between or within repositories. These records can be difficult to discern particularly where single specimens have been assigned multiple, or worse, no unique identifiers, or where the same locality has been georeferenced independently by multiple institutions. We provide a custom function called *dupeSummary*, that iteratively searches occurrences for duplication using multiple column-sets. Users can choose to identify duplicates based on identifier columns, collection information, or both. They may also define any number of custom column sets by which to identify duplicates.

Some identifier columns might contain codes that are too simple. The function allows users to set thresholds to ignore those occurrences when checking the relevant columns for duplicates. For example, the *catalogNumber* might be “145” or “174a”, which could result in over-matching of duplicates. Users may set a characterThreshold and numberThreshold which would ignore codes which don’t pass both. Users may also set a numberOnlyThreshold, which will check codes above that threshold, irrespective of the characterThreshold. We use the defaults of two, three, and five, respectively. Hence, minimum passing codes could include “AG194” and “390174”. This can be entirely turned off by setting all values to zero or ignored for selected column sets using CustomComparisonsRAW.

The function identifies duplicates based on collection information where it iteratively compares user-defined sets of columns. The function first compares custom column sets and can then compare generic, but customisable, column sets. For our bee data, we used several steps to identify duplicates. Firstly, we compared (i) *catalogNumber* and *institutionCode* using CustomComparisonsRAW. Secondly, we identified duplicates based on the *scientificName* with the (ii) *catalogNumber* and *institutionCode*, (iii) *gbifID*, (iv) *occurrenceID*, (v) *recordId*, and (vi) *id* columns using CustomComparisons. Finally, we identified duplicates using the generic column sets of *decimalLatitude*, *decimalLongitude*, *scientificName*, *eventDate*, and *recordedBy* columns at the same time as the (vii) *catalogNumber* and (viii) *otherCatalogNumbers* columns.

Using the *scientificName* column with the identifier columns allows different species with the same identifier to be maintained. This is important where an event identifier (multiple specimens from one collection event) has been placed in the wrong column. For the occurrences where the taxonomy did not match (e.g., because of an ambiguous or incomplete identification) duplicates won’t be identified; in these cases they will be removed by *harmoniseR* (or another flag). At the same time, occurrences where species identifications have been updated in only some datasets will not be identified as duplicates. Users may choose their own input parameters while weighing the costs and benefits.

The function arranges the data by (i) a user-defined list of input sources (where the first data sources are preferred over later ones; i.e., GBIF > SCAN > iDigBio, etc.), then, (ii) completeness by user-defined columns, and (iii) by the summary column (to keep clean occurrences). For point three above, duplicate occurrences with more of these completeness columns will be preferred over those with fewer; i.e., the most-complete record is chosen. Where public and private data were duplicated, we gave preference to private data providers over the public data aggregators under the assumption that data providers have the most recent information. We also preferred manually cleaned occurrence records from Chesshire, et al. ^93^ over those sourced directly from data aggregators. All pairwise duplicates were clustered where they overlapped and a single best occurrence was kept using the above arrangement. This step flagged 7,568,016 (41%) occurrences. Most of these duplicates arose between data sources; however, within-source duplicates were quite prominent in SCAN (Fig. 2).

**Figure 2.**
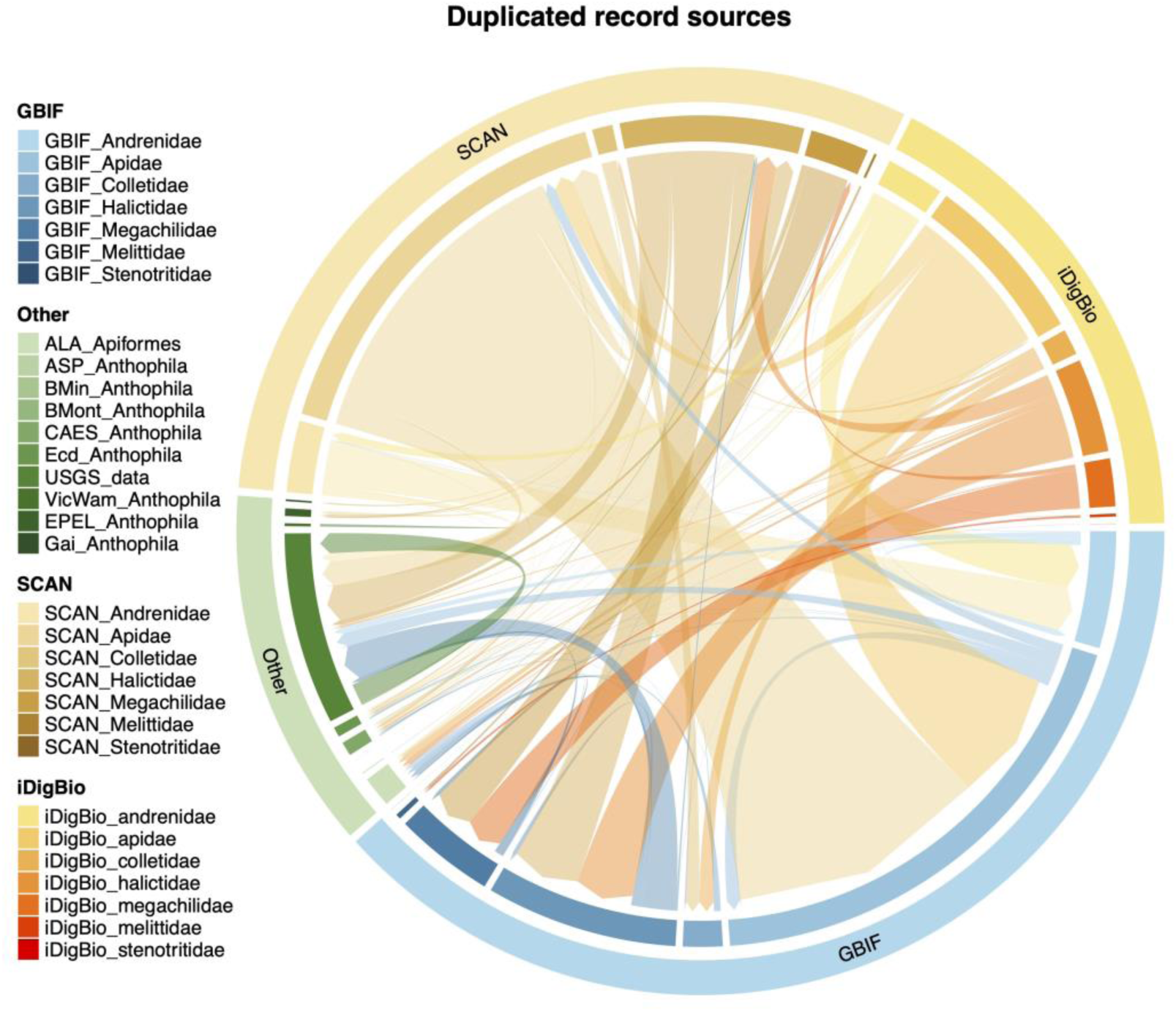
A chord diagram showing duplications between major data sources. The major data repositories, the Global Biodiversity Information Facility (GBIF), Integrated Digitized Biocollections (iDigBio), and Symbiota Collections of Arthropods Network (SCAN) are indicated on the diagram’s outer ring with bee families indicated on the associated inner ring. Smaller collections are indicated under ‘Other’ on the outer ring and the associated inner ring indicates individual datasets. The ‘Other’ collections are the Atlas of Living Australia (ALA), Allan Smith-Pardo (ASP), Bob Minckley (BMin), *Bombus* Montana (BMont), The Connecticut Agricultural Experiment Station (CAES^30,31^), Ecdysis (Ecd^28^), Gaiarsia (Gai^29^), Elinor Lichtenberg (Lic^37^), Victorian and Western Australian Museum (VicWam^5,39^), and the United States Geological Survey (USGS) data. The size of each ring and inner linkage (chord) is relative to the number of occurrences. The chords link occurrences are that are duplicated between data sources^87^.

### 8. Final filter

We first used the function *manualOutlierFindeR* function to flag occurrences identified as misidentifications or outliers by experts who reviewed interactive point maps. (from *interactiveMapR* below). There were 181 records from Chesshire, et al. ^93^ and 2,203 records identified by expert review of interactive species maps (see Technical validation). However, these were revised following the addition of new occurrence data. For each outlier identified, the function uses the output from *dupeSummary* to identify its duplicate records and flags those for removal. This function flagged 2,159 (<1%) records.

The “flagged-but-uncleaned” dataset was produced with the *summaryFun* which updated the .*summary* column. The .*summary* column is updated to include any records that are flagged in at least one flagging column, with user-defined exceptions. We excluded several filtering columns that we decided were not critical to data integrity. These were the diagonal and grid flags (*.sequential*, .*gridSummary*, .*lonFlag*, and .*latFlag*) and the taxonomic uncertainty flag (.*uncer_terms*). We also excluded high coordinate uncertainty (.*uncertaintyThreshold*) and records out of range (*.year_outOfRange*) because this filtering level might be too strict for general analysis and can remove ~1 million otherwise clean records at the 1 km level. Users may download our flagged dataset and begin at this step to choose the flags that are important for their hypotheses to remove or customise with different thresholds (OutputData/05_unCleaned_database.csv)^25^. For our “completely cleaned” dataset we followed used the above parameters in *summaryFun* but also chose to filter to only clean records and then remove all filtering columns (OutputData/05_cleaned_database.csv)^25^. This step removed 11,418,235 (62%) occurrences and left 6,890,148 (38%) cleaned occurrences.

### 9. Summary figures, tables, and outputs

Beyond the figures produced throughout the cleaning process by the *bdc* package, we also provide several unique custom figure functions.

#### 9.1 Duplicate chordDiagrams

We provide a function, *chordDiagramR*, that wraps the *circlize*^94^, *ComplexHeatmap*^95,96^, and *paletteer*^97^ packages to build a chord diagram that visualises the linkages between duplicated occurrence data sources (Fig. 2).

#### 9.2 Duplicate histogram

We provide a function, *dupePlotR*, to visualise duplicates by source, breaking them down into (i) discarded duplicates, (ii) kept duplicates (as chosen in *7.x Duplicate records*), and (iii) unique records (Fig. 3). This is displayed as both a total number of records and the proportion within each data source.

**Figure 3.**
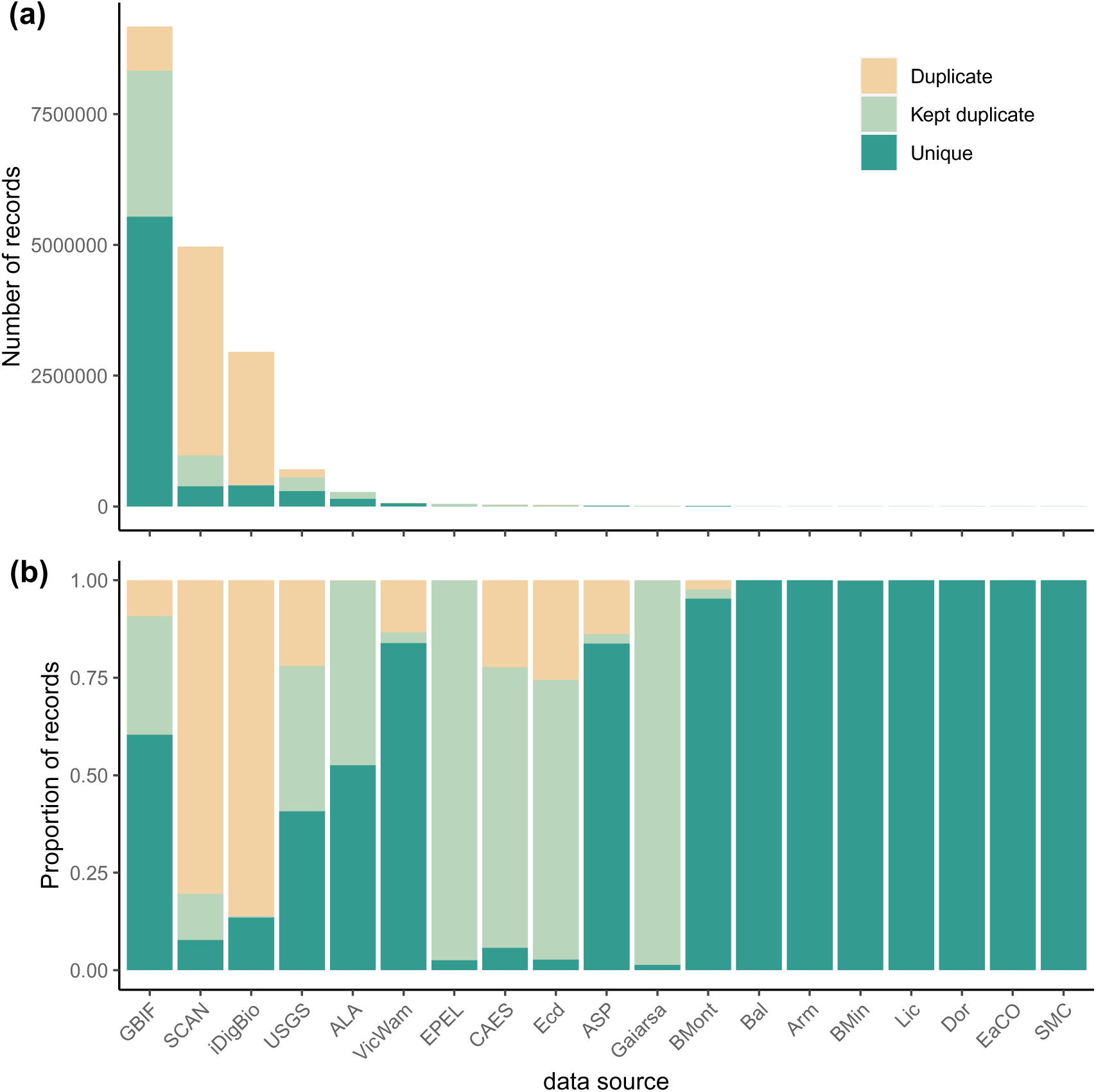
Duplicate occurrence summary. Two bar plots showing (a) the total number of records and (b) the proportion of records in each dataset that were duplicates (sand), kept duplicates (light green), and unique (dark green). Duplicates are occurrences that were identified to have a match in another or the same dataset and that were thus flagged or discarded. Kept duplicates are the same as duplicates, except they are the version of the occurrence records that were kept. Unique occurrences are those that were not matched to any other occurrences and were also kept. The included datasets are the Global Biodiversity Information Facility (GBIF), Symbiota Collections of Arthropods Network (SCAN), Integrated Digitized Biocollections (iDigBio), the United States Geological Survey (USGS), the Atlas of Living Australia (ALA), Victorian and Western Australian Museum (VicWam^5,39^), Elle Pollination Ecology Lab (EPEL^26^), the Connecticut Agricultural Experiment Station (CAES^30,31^), Ecdysis (Ecd^28^), Allan Smith-Pardo (ASP), Gaiarsia (Gai^29^), *Bombus* Montana (BMont^27^), Ballare, et al. ^36^ (Bal), Armando (Arm), Bob Minckley (BMin), Elinor Lichtenberg (Lic^37^), Eastern Colarado (EaCO), Texas literature data (SMC), and Dorey literature data (Dor^3,4,38^).

#### 9.3 Flags by source

We provide the *plotFlagSummary* function that produces a compound bar plot that, for each data source, indicates the proportion of records that pass or fail, or cannot be assessed for each flag (Fig. 4). We built additional functionality into this function that allows users to provide a species name (and the column in which that name is found) and (i) save the occurrence data, (ii) produce a map coloured by a filtering column of choice, and (iii) the compound bar plot for that specific species. This can be an excellent function to quickly examine potential issues. Additionally, users can choose to output the table used to make the figure using saveTable = TRUE.

**Figure 4.**
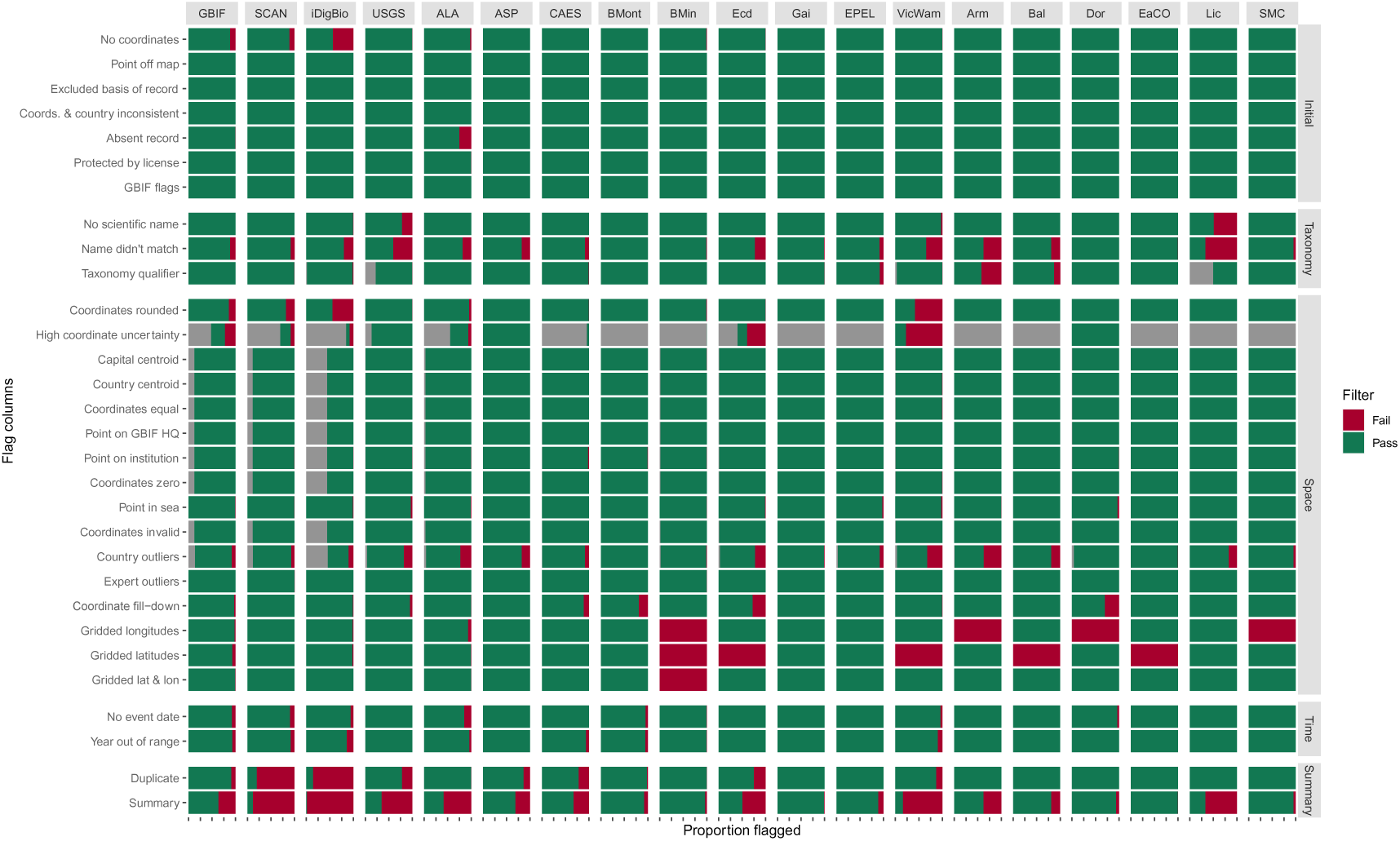
Flag summary. A compound bar plot showing the proportion of all occurrences within each data source (x-axis) that were flagged for each filtering step (y-axis). Green indicates the proportion of occurrences that passed a filter, red indicates the proportion that failed a filter, and grey indicates those that could not be assessed. Each x-axis tick indicates intervals of 25%. The summary row indicates the total proportion of passed or failed occurrences for each dataset based on those that were chosen to be filtered. In this instance, (i) taxonomic qualifier, (ii) gridded longitudes, (iii) gridded latitudes, and (iv) gridded latitude and longitude were not included in the summary row. The included datasets are the Global Biodiversity Information Facility (GBIF), Symbiota Collections of Arthropods Network (SCAN), Integrated Digitized Biocollections (iDigBio), the United States Geological Survey (USGS), the Atlas of Living Australia (ALA), Allan Smith-Pardo (ASP), the Connecticut Agricultural Experiment Station (CAES^30,31^), *Bombus* Montana (BMont^27^), Bob Minckley (BMin), Ecdysis (Ecd^28^), Gaiarsia (Gai^29^), Elle Pollination Ecology Lab (EPEL^26^), Victorian and Western Australian Museum (VicWam^5,39^), Armando (Arm), Ballare, et al. ^36^ (Bal), Eastern Colarado (EaCO), Elinor Lichtenberg (Lic^37^), Texas literature data (SMC), and Dorey literature data (Dor^3,4,38^).

#### 9.4 Maps

We provide a function, *summaryMaps*, that uses the cleaned data to show the number of species and number of occurrences per country (Fig. 5). For the latter function, we broke up these values using classes and “fisher” intervals^98^; however, any style from the *classInt*^99^ package can be entered. Additionally, we provide a function, *interactiveMapR*, that iteratively makes and saves interactive html maps for any number of species. These interactive maps colour occurrences if they pass or fail any flags, if they are country outliers, or if they are expert outliers. Collection and flag information are provided in pop-ups for each point.

**Figure 5.**
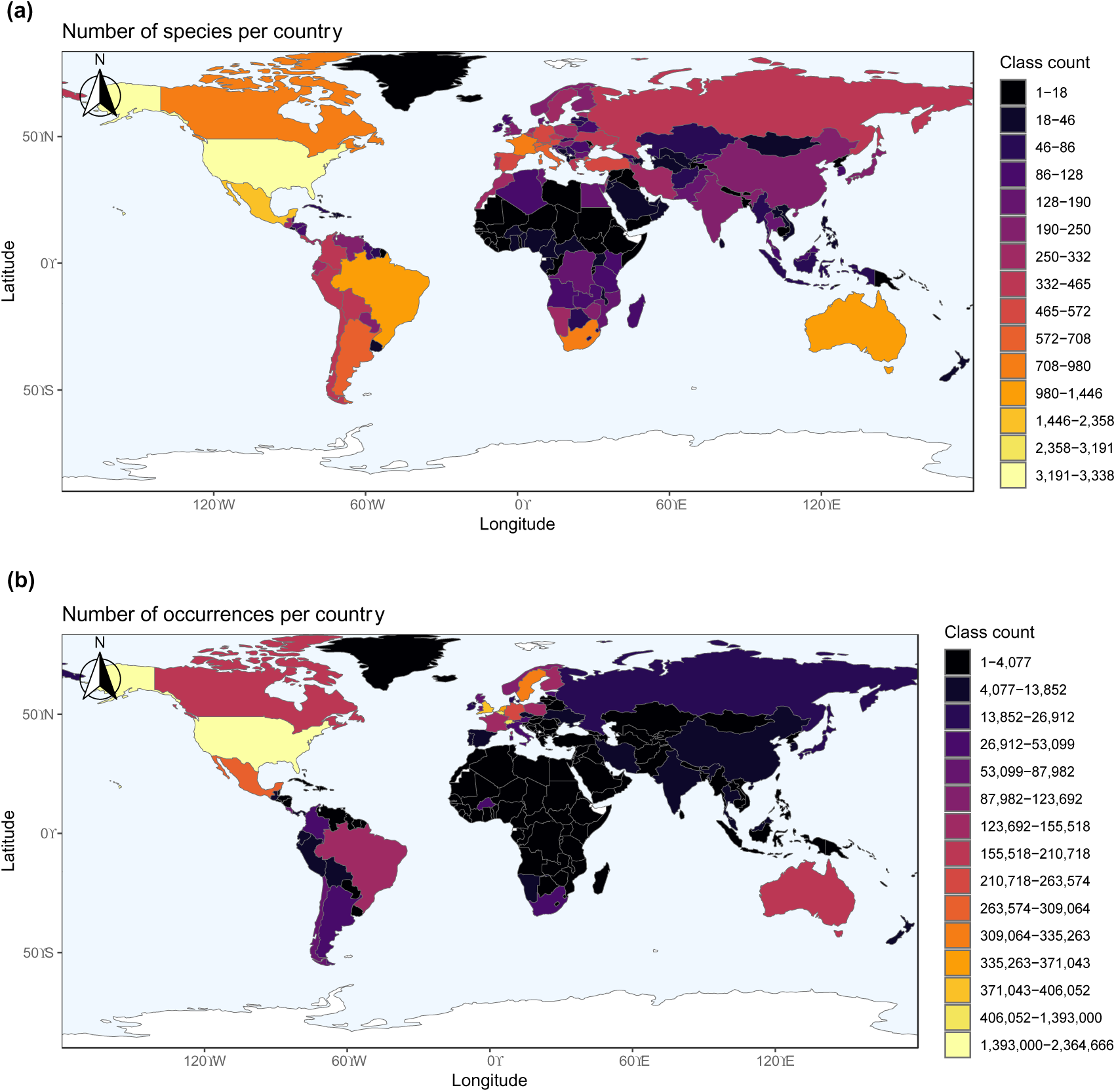
Occurrence-country summary maps created using the cleaned data indicating the (a) number of species per country and (b) number of occurrences per country from the filtered data. Colours indicate the number of (a) species or (b) occurrences where dark colours are low and yellow colours are high. Class intervals were defined using a “fisher” method.

#### 9.5 Data providers

We provide a function, *dataProvTables*, that produces a table of data providers with the number of cleaned occurrences and species for each. The function also attempts to identify the data provider using other columns when it is omitted from the *institutionCode* column, providing an updated institution code and name. This function is thus far customised to bee datasets. We provide both the top 14 rows (for datasets with >100,000 clean occurrences; Table 1) and the full table (OutputData/Reports/dataProviders.xlsx)^25^.

**Table 1.** The institution code, institution name, number of clean bee occurrences, and number of unique bee species for the datasets with >100,000 clean bee occurrences in our dataset.

| Institution code | Institution name | Occurrence count | Species count |
| --- | --- | --- | --- |
| iNaturalist | iNaturalist | 827,600 | 2,952 |
| USDA-ARS | USDA Agricultural Research Service | 451,180 | 4,272 |
| Observation.org | Observation.org | 421,201 | 691 |
| SLU Artdatabanken | SLU Artdatabanken | 355,440 | 291 |
| Natuurpunt | Natuurpunt | 268,470 | 303 |
| USGS | United States Geological Survey Bee Lab | 254,173 | 1,247 |
| CSCF | Swiss National Biodiversity Data and Information Centres | 246,684 | 578 |
| AMNH | American Museum of Natural History | 244,432 | 5,153 |
| KU | Kansas University Entomology collection | 218,045 | 5,765 |
| Biological Records Centre | Biological Records Centre | 191,790 | 258 |
| Bumblebee Conservation Trust | Bumblebee Conservation Trust | 148,489 | 273 |
| AMU | Adam Mickiewicz University in Poznań | 144,805 | 191 |
| naturgucker | naturgucker | 139,950 | 411 |
| FinBIF | Finnish Biodiversity Information Facility | 135,463 | 315 |

#### 9.6 Flag summary

We built a function, *flagSummaryTable*, that will produce a summary table of the total number of failed occurrences per flag and per species (or any other categorical column of interest).

#### 9.7 Taxonomic and country checklist queries

Users may be interested in querying *BeeBDC* for species names, validities, and in which countries a species can be found. Our function, *BeeBDCQuery*, allows users to enter one or more species name as a character vector and it will report on (i) the validity of the name, (ii) all synonyms, and (iii) the countries in which it is found. This information will be returned as a list of tables.

### 10. Package data

#### 10.1 Bee taxonomy and checklist

We provide a modified version of the global bee taxonomy and country-level checklist produced by John Ascher and John Pickering and hosted on Discover Life^23^ as part of *BeeBDC*. These datasets are downloadable directly from the figshare using the functions *beesTaxonomy* and *beesChecklist*, respectively.

#### 10.2 Test datasets

We further provide three test datasets with BeeBDC. The dataset *bees3sp* includes 105 flagged points from three haphazardly chosen species; *beesFlagged* contains 100 randomly chosen flagged occurrence records; and *beesRaw* contains 100 randomly chosen unflagged occurrence records.

## Data Records

Our dataset and the data used in our analyses are available for download in our figshare repository (https://doi.org/10.25451/flinders.21709757)^25^. See Data Sources above for further information. Occurrence datasets are provided in standard DarwinCore format and saved as .csv files. The additional materials are organised into the following folders:

1. Figures. Contains primary figures and additional *bdc* figures.
2. OutputData. Contains the “cleaned” (05_cleaned_database.csv) and “flagged-but-uncleaned” (05_unCleaned_database.csv) datasets, reports, and the *R* console outputs from the script (RunNotes_BeeBDC_1Sep23.txt).
3. InputData. Contains the major repository downloads, additional input datasets, and custom Chesshire, et al. ^93^ data files.
4. ExtraTables. Contains metadata for the major repository downloads (MajorRepoAttributes_2023-09-01.xlsx) and the added taxonomy variants (AddedTaxonomyVariants.xlsx).
5. The beesTaxonomy.Rda and beesChecklist.Rda datasets that are downloaded using BeeBDC (see 10. Package data above).

## Technical Validation

For our current data version, we outline the data quality according to our flags. We provide summaries of the deduplication process, including the duplication linkages within and between datasets (Fig. 2), and the number and proportion of duplicates, kept duplicates, and unique occurrences per dataset (Fig. 3). Importantly, we provide a figure summary of the proportion of records flagged for all data sources (Fig. 4). This figure indicates data quality across the entire dataset and where each data source might focus their cleaning efforts. We demonstrate the country-level patterns of both species and occurrences on a global map (Fig. 5). Finally, all *BeeBDC* and *bdc* summary and data quality figures and maps are provided in the Additional Figures folder (https://doi.org/10.25451/flinders.21709757).

We assessed our figures and interactive maps in a lengthy and iterative error-checking process. Throughout the development of the package, functions and results were scrutinised by the authors, particularly JBD, EEF, AN-B, RLO’R, SB, DAG, DdP, K-LJH, LMM, TG, TAZ, MCO, LMG, JSA, ACH, and NSC. Functions were tested throughout development and as a part of our CRAN submission process to ensure robustness and consistency of results with expectations. Occurrence records were examined frequently in this process and, several times, interactive maps of 100 randomly chosen species were produced and manually checked. In addition to this, interactive maps for all species with sufficient data between Colombia and Canada (4,221 spp.) were produced and manually checked (in some cases for multiple versions) as part of an additional effort to build species distribution models of the bees in that region led by AN-B and NSC.

We found that data quality was highly variable between sources (Fig. 4). Data duplication was often the most prominent flag, comprising 41% of the total uncleaned dataset. There was substantial duplication between and even within both major and minor data repositories (Figs. 2 and 3). Importantly, the assumption that all data make their way to GBIF is incorrect (at least at the time of downloading); we found that each repository holds unique data (Fig. 3). However, it is also likely that poor data quality might preclude successful duplicate matching. Regardless, this highlights the importance of sourcing occurrence data from a variety of repositories. However, in terms of actual input data quality and completeness, many occurrences (i) lacked coordinates, (ii) did not have a scientific name or that name did not match known taxonomy, (iii) had low-resolution coordinates, (iv) had high coordinate uncertainty (where this was estimated), (v) did not match the country checklist on the Discover Life website, (vi) lacked collection date, and/or (vii) were collected prior to 1950 (Fig. 4). These quality issues might preclude data use in downstream works and require further efforts to repair. In worst case scenarios, unfit data might pass flagging steps and cause misleading results. The presented workflow enables researchers to prepare data that can support robust analyses. To the best of our knowledge, this is now the most thoroughly curated global occurrence dataset for bees and is potentially leading for any terrestrial invertebrate group of this size.

The mismatch between bee species richness according to occurrence data and country checklists has already been highlighted by Orr, et al. ^6^. We also highlight the geographical, taxonomic, and collection biases in the global bee data (Fig. 5). We show that the number of cleaned species occurrences in most of Africa and large parts of Asia are in the lowest class counts, where classes are defined using a Fisher-Jenks algorithm^98^ (Fig. 5a). We also show that the number of occurrences in Africa, Asia, and even South America are often in the lowest class counts and to a greater degree than species counts (Fig. 5b). Some countries in Africa (Equatorial Guinea, Djibouti, and parts of Western Sahara) do not have a single cleaned occurrence record (Fig. 5). There are also large mismatches between the number of species and number of occurrence records. These disparities are hugely concerning and highlight ongoing inequalities in taxonomic, sampling, and digitisation efforts. While we integrate new datasets from Central and South America (with more expected to be released in future versions) we note that there are likely many more data available in grey literature for the under-sampled regions of the world and museum specimens that are yet to be digitised. We hope that this contribution provides a foundation for a more formal recognition and prioritization of important taxonomic, sampling, and digitisation efforts.

## Usage Notes

All of our occurrence datasets are cleaned for (i) taxonomic information according to the Discover Life website, (ii) country name (country_suggested) and ISO2 code, and (iii) recoverable event dates from other columns. When using our data or parts of our workflow users should also cite the data sources where they are relevant. Our occurrence data are provided in two ways:

### Cleaned

We provide a dataset that is completely cleaned of records that failed any of our filtering steps with the exception of: (i) .*gridSummary,* (ii) .*lonFlag*, (iii) .*latFlag*, (iv) .*uncertaintyThreshold*, (v) .*sequential*, and (vi) .*year_outOfRange*. All filtering columns are also removed from this dataset to reduce file size.

### Flagged-but-uncleaned

We provide a dataset that includes all of the above filters as flags, in addition to the otherwise unfiltered flags above. This dataset is provided with a column for each of these flags where occurrences that failed for a filtering step are flagged as “FALSE” and those that passed as “TRUE”. This convention is used to maintain continuity with functions from the *bdc* and *coordinateCleaner* packages. Users are able to use our script to read in and filter their data or manually filter these data in a way that is appropriate for their use. Our filtering script should also provide users with the necessary template to reduce or modify their data into their desired size and structure. In particular, users may use the dontFilterThese argument in the *summaryFun* function to exclude certain filters that do not relate to their research question.

Users may incorporate our data pulls or conduct their own. However, we do note that ALA, through the *galah* package, is the only major data provider with a convenient download protocol for large datasets. Other Application Programming Interfaces (APIs) and even web protocols make downloading a limiting and lengthy process; for example, the *rgbif* package has a limit of 100,000 occurrences. Making similar API methods available for other major data repositories would greatly increase their accessibility. Usage of the *BeeBDC* package and workflow should be accessible to most users with basic *R* knowledge. Once the datasets and folder structure are determined the script should require minimal user input. We highlight again that this workflow was tested on a machine with 64 GB of RAM and that users with less RAM might have prolonged run times and need to reduce chunk sizes for some functions. In the end, this work is aimed to facilitate open source data for both scientific and applied usage.

## Code Availability

We provide a full website with vignettes (including a complete and a basic workflow) that demonstrates the functionality of our script with the input data (https://jbdorey.github.io/BeeBDC/index.html). Our “basic workflow” is for users that simply wish to download our flagged but unfiltered dataset and apply (i) manual filters based on our filtering columns or further filter to only include certain (ii) countries, (iii) date ranges, or (iv) uncertainty levels. Secondly, we provide these annotated *R-*scripts run from start to finish on our GitHub (https://github.com/jbdorey/BeeBDC/tree/main/inst). Our scripts, related files, and downloaded instructions can be found on our GitHub page (https://github.com/jbdorey/BeeBDC).

## Author contributions

The project was **conceptualised** by JB Dorey and KLJ Hung. The project design was guided by JB Dorey, EE Fischer, MS Rogan, YV Sica, MC Orr, LM Guzman, JS Ascher, AC Hughes, and NS Cobb. **Data were provided and/or curated** by JB Dorey, PR Chesshire, A Nava-Bolaños, SM Collins, EM Lichtenberg, EM Tucker, A Smith-Pardo, A Falcon-Brindis, DE de Pedro, KA Parys, RL Minckley, T Griswold, T Zarillo, JA Ascher, AC Hughes, and NS Cobb. The **code was drafted** by JB Dorey, tested by JB Dorey and EE Fischer, and input was provided by RL O’Reilly, S Bossert, and SM Collins. *R* Markdown was generated by JB Dorey and R O’Reilly. Figures were created by JB Dorey and S Bossert. The **manuscript was drafted** by JB Dorey and all authors provided critical feedback and reviewed the final manuscript. First and senior authors are ordered by relative contribution and alphabetically in between these groups.

## Acknowledgements

We acknowledge the wonderful data providers such as ALA, GBIF, SCAN, and iDigBio, who maintain and make available datasets for public use. We also sincerely thank Sam Droege for organising and contributing the substantial USGS bee dataset. We additionally acknowledge and thank individuals and organisations that do the same, including those individually used here produced by: Elizabeth Elle, Casey Delphia, Ecdysis, Gaiarsa et al., Arathi Seshadri, and the Florida State Collection of Arthropods. A special thanks also goes to Elizabeth Murray for her excellent suggestion that led to the naming of our R package, BeeBDC. Mention of trade names or commercial products in this publication is solely for the purpose of providing specific information and does not imply recommendation or endorsement by the U.S. Department of Agriculture. USDA is an equal opportunity provider and employer. JB Dorey was partly funded by the Biodiversity Outreach Network and at the earliest stages of the paper’s conception was partly funded by Burt’s Bees. EE Fischer was supported by the Hans Rausing Scholarship in the History of Science. A Nava-Bolaños is grateful to Consejo Nacional de Ciencia y Tecnología for the postdoctoral fellow for the project “Polinizadores: actores clave en la seguridad alimentaria”. S Bossert was funded under NSF grant, DEB-2127744. EM Lichtenberg and SM Collins were partially funded through the Texas State Wildlife Grants program grant CA-0002506 in cooperation with the U.S. Fish and Wildlife Service, Wildlife and Sport Fish Restoration Program. MS Rogan, YV Sica, and W Jetz were partly funded by Burt’s Bees. TA Zarrillo was additionally funded by Hatch funds from the Connecticut Agricultural Experiment Station and the Connecticut Department of Energy and Environmental Protection Wildlife Division and the federal State Wildlife Grants Program. Additional partial funding was provided by the iDigBees TCN NSF award #2216927.

## Tables

**Table S1.** The occurrence data filtering steps, functions, packages (*BeeBDC*^24^, *bdc*^13^, and *CoordinateCleaner*^76^), descriptions, and function references required to undertake each step of filtering. The function steps include script and data preparation, flagging (adding function result column), carpentry (altering the data itself), and filtering (removing occurrences based on data flags). The packages *BeeBDC* and bdc also have a selection of functions that are useful for data visualisation and table production which are critical for error-checking the results. Parentheses under descriptions often indicate the flagging columns produced by that function. Asterisks next to function names indicate that they are parallel-ready (can be run across multiple threads).

